# JLOH: Inferring Loss of Heterozygosity Blocks from Sequencing Data

**DOI:** 10.1101/2023.05.04.539368

**Authors:** Matteo Schiavinato, Valentina del Olmo, Victor Njenga Muya, Toni Gabaldón

**Affiliations:** Barcelona Supercomputing Centre (BSC-CNS). Plaça Eusebi Güell, 1-3 08034 Barcelona, Spain; Institute for Research in Biomedicine (IRB Barcelona), The Barcelona Institute of Science and Technology, Baldiri Reixac, 10, 08028 Barcelona, Spain; Current address: Eskişehir Osmangazi University, Büyükdere Meşelik Yerleşkesi, 26040 Odunpazarı/Eskişehir, Turkey; Catalan Institution for Research and Advanced Studies (ICREA), Barcelona, Spain; Centro de Investigación Biomédica En Red de Enfermedades Infecciosas (CIBERINFEC), Barcelona, Spain

**Keywords:** Heterozygosity, LOH, hybrids, variation, sequencing, pipeline

## Abstract

Heterozygosity is a genetic condition in which two or more alleles are found at a genomic locus. Among the organisms that are more prone to heterozygosity are hybrids, i.e. organisms that are the offspring of genetically divergent yet still interfertile individuals. One of the most studied aspects is the loss of heterozygosity (LOH) within genomes, where multi-allelic sites lose one of their two alleles by converting it to the other, or by remaining hemizygous at that site. LOH is deeply interconnected with adaptation, especially in hybrids, but the *in silico* techniques to infer LOH blocks are hardly standardized, and a general tool to infer and analyse them in most genomic contexts and species is missing. Here, we present JLOH, a computational toolkit for the inference and exploration of LOH blocks which only requires commonly available genomic data as input. Starting from mapped reads, called variants and a reference genome sequence, JLOH infers candidate LOH blocks based on single-nucleotide polymorphism density (SNPs/kbp) and read coverage per position. If working with a hybrid organism of known parentals, JLOH is also able to assign each LOH block to its subgenome of origin.

## 1. Introduction

Heterozygosity is the genetic condition that exists when two or more alleles are present at a specific genomic locus. Genomes with ploidy higher than one can exhibit different degrees of genomic heterozygosity, which is generally described as the amount of heterozygous sites across their sequence. Genomic heterozygosity entails genetic diversity, and a wider pool of alleles upon which selection can act. Heterozygosity is widespread and pervades all non-haploid organisms, but is particularly high and relevant for hybrid organisms. Hybrids are either the offspring of two diverged organisms from different species (inter-species hybrids) or from genetically distant populations of the same species (intra-species hybrids) (1). Hybridisation is pervasive across the eukaryotic tree of life, and is at the origin of many lineages of fungi, plants, and other eukaryotes (2). Hybrids carry a chimeric genome, resulting from the combination of divergent chromosomes from the parental organisms, i.e. homeologous chromosomes (3). As a result, non-haploid hybrids are highly heterozygous, as homeologous chromosome loci can present different alleles inherited from each of the parents. The level of heterozygosity will depend on the initial divergence of the parental lineages, and will subsequently evolve by processes that include differential loss of chromosomes and recombination between homeologous genomic regions (4–7). Some of these processes lead to loss of heterozygosity (LOH) resulting in particular genomic patterns in which homozygous and heterozygous segments (called blocks) alternate along a chromosome (8). Differences in LOH patterns are often the main source of variability among individuals of hybrid populations or species, and they can be the main drivers of phenotypic differences (7). Hence, there is a growing interest in accurately inferring LOH patterns in population genomics studies. In recent years, many studies have focused on studying LOH in relation to the biology and evolution of hybrids (7, 9), but also in non-hybrid contexts such as cancer (10), and niche adaptation (11). One area of current research is the evolution of hybrid lineages such as those of the yeast genus *Saccharomyces* (12). The *Saccharomyces* genus harbors eight *sensu stricto* species which can all hybridize between each other, resulting in hybrids with different levels of parental genetic divergence. *Saccharomyces* hybrids have been documented, which carry a subgenomic sequence divergence of 25% or more (12), underscoring the ability of these species to hybridize across large evolutionary distances. In the past decade, many studies have focused on inferring LOH patterns from genomes of yeast hybrids, resulting in novel insights on hybrid genome evolution that facilitate comparative analyses (7, 9, 13–17). Most approaches to infer LOH patterns at the genome-wide level are based on mapping short read sequencing data to a genome reference (18–20). One of the earliest pipelines is YMAP (21). This pipeline was designed to analyse variation data from the human pathogen *Candida albicans*, taking as input microarray, high-throughput sequencing or RAD-Seq data. Among other things, YMAP would return an LOH profile extracted from the SNP profiles deriving from read mapping. Generally, LOH information is extracted from the combination of SNP data and read coverage data. However, these data are most often analysed semi-manually, using ad-hoc pipelines and parameters which are hardly reproducible in other environments. An example of this is a recently published pipeline named MuloYDH (22). MuloYDH is designed to analyse the variation landscape in hybrid yeast genomes, inferring also LOH blocks from variation data. The pipeline includes many custom scripts wrapped into a bash script with few tunable arguments, therefore limiting the customization of analysis. A few programs exist to infer LOH tracts in cancer setups (23, 24), but a general, easy-to-use tool that requires only a few common input data types, is not designed for a specific use-case, and that can handle hybrid genomes is missing. Considering that the study of LOH is gaining momentum, we undertook a more general approach, reducing the input requirements to the bare minimum and implementing a scalable computational solution that returns reproducible results in a highly-parallelized fashion with easy parameter customization. We imagined the typical use-case of a general LOH inference software and implemented a toolkit named “JLOH”, which performs a series of steps to infer and analyse LOH blocks starting from mapped reads, single-nucleotide polymorphisms (SNPs) and a reference genome sequence. These three input data types are highly common in genomic analyses, therefore allowing those who are already performing variation studies to add one extra layer of information to their workflow (i.e. LOH blocks) without generating new files or having to deal with cumbersome pipeline setup processes. JLOH uses SNP density to define heterozygous and homozygous regions of the provided reference genome sequence, and from those regions infers candidate LOH blocks using read coverage information as a means of validation. JLOH has been especially (although not exclusively) designed to work in the context of hybrid species, allowing researchers to disentangle LOH blocks in these complex genomes. We first test JLOH’s ability to infer LOH blocks from simulated read data that is progressively more divergent from the reference genome onto which they are mapped. We then test its performance on public sequencing data from *Saccharomyces* hybrids (18) and from the pathogenic hybrid species *Candida orthopsilosis*. Our results show that JLOH is reliable in inferring LOH blocks in all these settings with great accuracy.

## 2. Materials & Methods

### 2.1 Code implementation for LOH block inference

The main module of JLOH is “jloh extract”, which infers LOH blocks from three input files: a reference genome sequence, a BAM file with reads mapped onto this genome, and a VCF file containing single nucleotide polymorphisms (SNPs) that were called from these mapped reads. The LOH block inference workflow performed by “jloh extract”, is represented in Figure 1.

**Figure 1.**
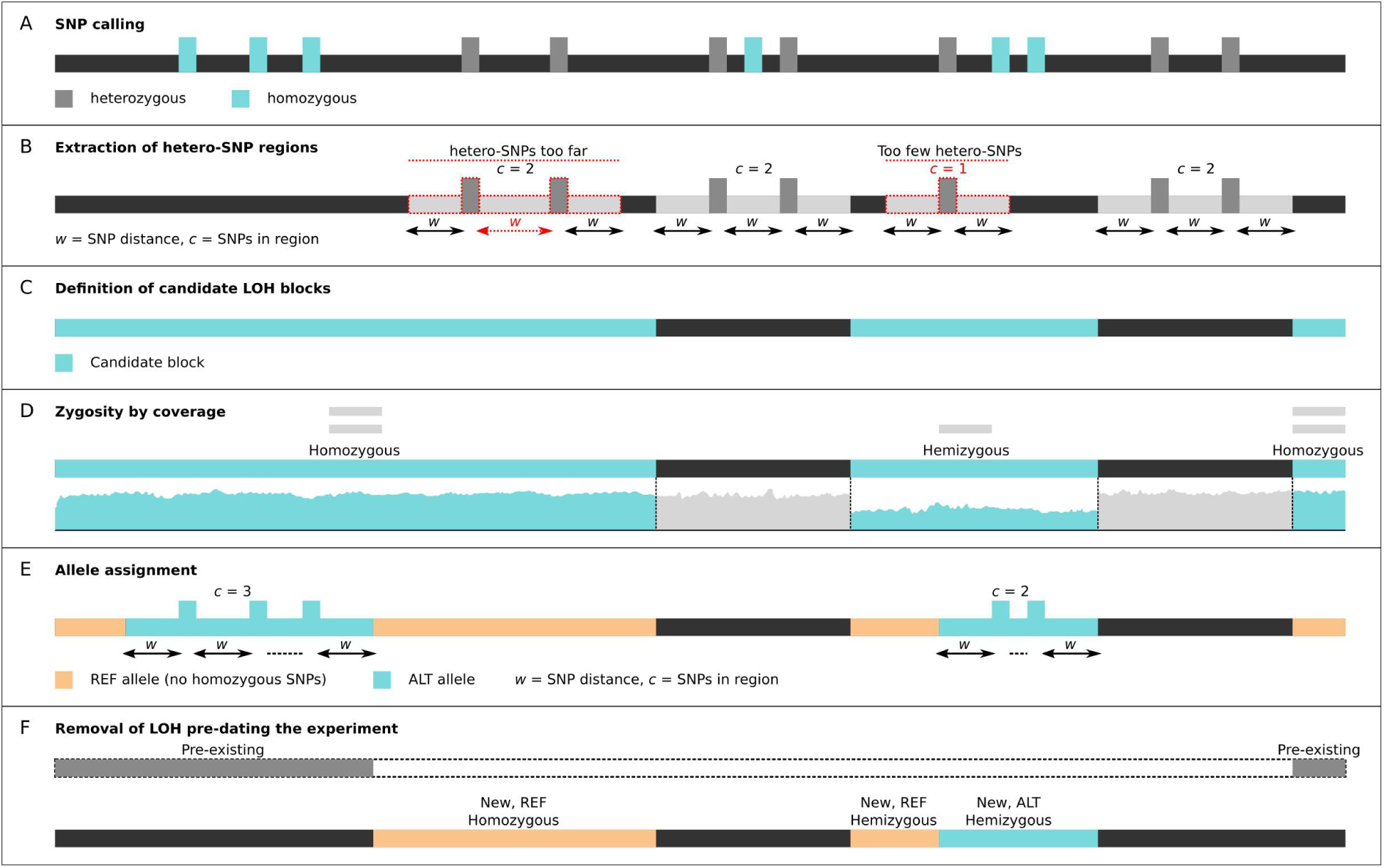
Schematic representing the rationale of “jloh extract”. **A**) SNPs are parsed by the program from the provided VCF file(s), subdividing them into heterozygous (gray) and homozygous (cyan). **B**) Regions rich in heterozygous SNPs are identified (light gray) based on SNP density criteria. Dotted red lines represent intervals containing heterozygous SNPs that do not fit the SNP density thresholds either due to the number of SNPs (c) or to the distance between them (w). **C**) Candidate LOH blocks, i.e. regions depleted in heterozygous SNPs are extracted (cyan), which are complementary to the identified heterozygous regions. **D**) Coverage per position is assessed for each candidate LOH block (cyan) and zygosity is inferred from it (horizontal light gray bars). **E**) When using data from hybrids, homozygous SNPs are used to subdivide candidate LOH blocks into reference (REF, orange) and alternative (ALT, cyan) allele based on the same mechanism explained in (B) for heterozygous SNPs. **F**) If specified by the user, candidate LOH blocks overlapping known LOH blocks are filtered out, leaving only the newly found ones.

SNPs provided as input are separated by the program in homozygous and heterozygous based on their genotype annotation (0/0, 0/1, 1/1, 1/2). In addition, to make the selection more stringent, the user can define a range of allele frequencies (“AF” annotation in VCF format) in which a SNP is considered heterozygous using the --min-af and --max-af parameters, both ranging from 0 to 1. Both SNP groups are clustered by reciprocal distance as long as they form a cluster with a SNP density (SNPs/kbp) greater or equal to the one passed via the --min-snps-kbp parameter. This parameter is the most impactful of the program (see results). By passing two comma-separated integer values, it allows the user to set a different SNP/kbp threshold for heterozygous and homozygous SNPs (e.g. --min-snps-kbp 15,3), which is key to eliminate many false positives in the final results (see benchmark results below). Genome intervals defined by clusters of heterozygous SNPs are assumed to be intervals that do not contain candidate LOH blocks, as their levels of heterozygosity are above the threshold. The complementary ones, which have a heterozygous SNP density below the threshold, are the raw candidate LOH blocks and are screened further. First, they are screened for the presence of homozygous SNPs. Those that are defined by a density of homozygous SNPs above the threshold are assumed to be candidate blocks that retained a haplotype different from the one of the reference genome. The ones that have a density of homozygous SNPs below the threshold, instead, are assumed to carry the same haplotype of the reference genome. Overlaps between heterozygous intervals and the candidate LOH blocks, which may arise due to the SNP distance clustering strategy, are trimmed down from the candidate blocks. Candidate LOH blocks that are longer than a certain size (adjustable with the --coarseness parameter) are retained. The surviving candidate LOH blocks are inspected further using coverage information from the BAM file. The read coverage per position of each candidate is computed, and those candidates where the fraction of covered positions falls below the one declared with the --min-frac-cov parameter are discarded. Note that, in LOH blocks, one of the two original alleles has to be present, hence the removal of totally uncovered regions. Coverage information is then used to compare the block to its surroundings. The block’s coverage is compared with the region upstream and downstream of it, the length of which can be adjusted through the --overhang parameter. If the block has substantially lower coverage than its surroundings, it is considered as an LOH event where one copy of the genome portion was lost and the remaining one is now hemizygous. The coverage ratio with the surroundings at which this is triggered can be adjusted through the --hemi parameter (e.g. --hemi 0.75). Any block that does not fit the hemizygous coverage threshold is considered homozygous. The LOH blocks that survive this process are reported both in a BED file and in a more human-readable tab-separated format (TSV) file, the latter containing all the necessary information about each block, such as the mean coverage per position, the allele (REF or ALT), the zygosity (homo or hemi), and the number of heterozygous and homozygous SNPs contained.

### 2.2 Inferring LOH subgenomic origin in hybrids with known parentals

The most relevant feature of the “extract” module is its ability to assign a subgenomic allele to each detected LOH block when working with a hybrid species that has known parentals. This is activated using the --assign-blocks parameter. This mode runs the same algorithm twice, once per parental genome. Note that the large majority of hybrids come from unknown parents: in that case, this assignment is not possible and the user may use the default “jloh extract” mode, still being able to retrieve LOH blocks although without a hint at which parent was favoured. The --assign-blocks mode requires two BAMs, two VCFs, and two reference genomes, one per parental species of the hybrid. The BAMs and the VCFs must be derived from mapping the reads sequenced from the hybrid onto the two parental genomes separately, and with relaxed parameters that allow crossmapping between subgenomes. This is achieved in the read mapping step by allowing twice as many mismatches between reads and reference than the actual genome divergence between subgenomes. If unsure about this parameter, the user may also use the “sim” module to simulate a genome at a certain divergence level and perform a mock run with those at different mapping stringency criteria; an example on how to use this module is provided in the “sensitivity and specificity” section. Through this crossmapping, differences between subgenomes will be highlighted as heterozygous SNPs and will be handled accordingly by the “extract” module. In this way, the inferred LOH blocks will represent genomic regions where the hybrid has lost inter-subgenomic diversity, losing one of the subgenomic alleles but retaining the other.

### 2.3 Implementation of other modules

JLOH is a toolkit composed of multiple modules that can be used to infer, filter and analyse LOH blocks (Figure 2). The most important of them is “jloh stats”. In a non-simulated scenario, the density and distribution of hetero- and homozygous variants across the genome can be highly variable. Hence, the optimal --min-snps-kbp parameter setting to pass to “jloh extract” might vary between individual samples, let alone different species, and it may be difficult for an inexperienced user to set this parameter. To account for this, the “jloh stats” module computes the mean density of homo- and heterozygous variants across the genome in windows of adjustable size. It detects and reports to the user the SNP density distribution quantiles which can be used to set the --min-snps-kbp parameter in an informed way, under the assumption that LOH blocks will be found at low heterozygous SNP densities and variable homozygous SNP densities.

**Figure 2.**
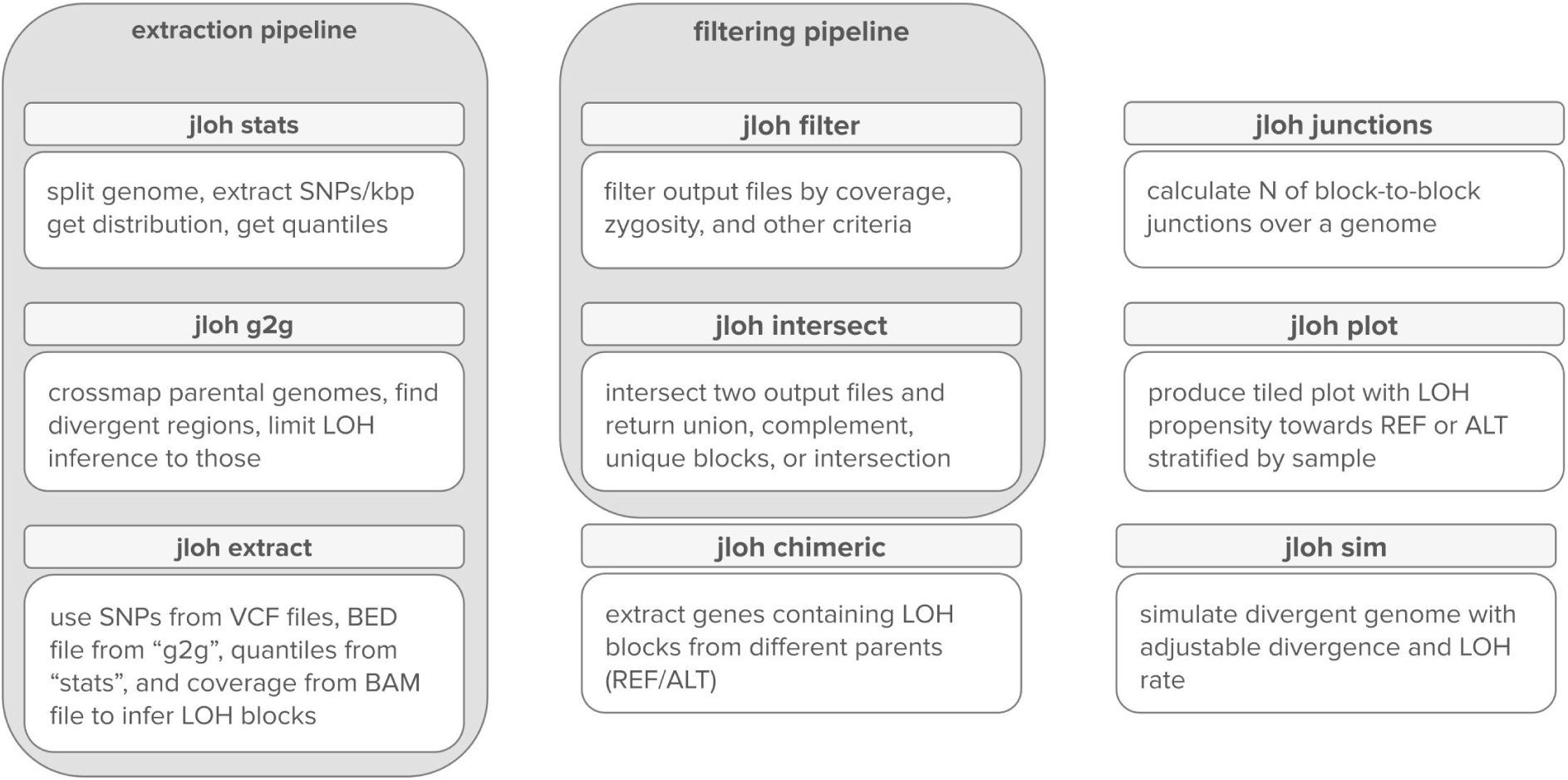
Schematic displaying the different modules available within the JLOH toolkit, with brief explanation.

The heterozygous SNP density Q10 quantile, for example, indicates a heterozygous SNP density which only 10% of the computed windows do not reach. In other words, 10% of the windows have a lower heterozygous SNP density than the Q10 heterozygous quantile. The same rationale is applied to homozygous SNPs. The “jloh stats” module produces a table with all the relevant quantiles, and the user is encouraged to choose among them based on the expected divergence between their reads and their reference genome. If, for example, they expect their reads to be 5% divergent (5 SNPs every 100 bp) from the reference genome, they may expect a mean heterozygous SNP density of ∼50 SNPs/kbp. This value should be close to the one reported a 50th quantile (Q50). The user may, therefore, expect candidate LOH blocks to retain some heterozygous SNPs in their sequence, although at a lower density than 50 SNPs/kbp. By passing to the --min-snps-kbp the SNP densities described in the 5th quantile (Q5), the user will retain most of the candidate blocks. By using upper quantiles, the user will be more stringent in the selection based on SNP density. As shown in the results (*C. orthopsilosis* section) this choice may be crucial.

A second relevant module is “jloh g2g”. Specifically when using hybrid genomes, there is a possibility that the two parental genomes may have regions of high homology. These regions are likely to produce a very small number of SNPs, hence looking like LOH blocks to JLOH. However, since they are already very similar at the parental level, these regions must not be considered as candidate LOH blocks. To take this into account, JLOH has a module named “g2g” (“genome to genome”). “jloh g2g” leverages the nucmer mapping tool from the MUMmer alignment software (25) to align the two provided parental genomes onto each other, and highlight regions that contain differences. These are the regions where JLOH has power to detect LOH, while those that carry no difference at all do not allow LOH detection given the absence of heterozygous SNPs. Note that, without this selection, these regions will be producing artifacts due to the lack of heterozygous SNPs and presence of read coverage. The produced BED file can be passed directly to “jloh extract” as argument for the --regions parameter. Alternatively, the user may pass a BED file with any region of interest, and “jloh extract” will return only the blocks that overlap these regions.

### 2.4 Simulated data generation

To test the validity of the “jloh extract” algorithm, a series of hybrid genomes with known LOH blocks were simulated starting from the *S. cerevisiae* S288C genome (26). The R64-1-1 assembly of the S288C strain of *S. cerevisiae* was used for the generation of simulated data. In total, 24 diverging versions of the *S. cerevisiae* genome were simulated, each with a different genetic divergence (1%, 3%, 5%,10%, 15%, 20%) and rate of loss of heterozygosity (10%, 20%, 30%, 40%). These operations were carried on with the “jloh sim” module. Note that the “sim” and the “extract” modules are completely independent from each other, and therefore cannot influence each other’s results (i.e. “extract” does not “know” the true positives generated by “sim”). This module breaks the genome sequence into arbitrary haplotypes of average size defined by the --mean-haplotype-size parameter. The size of each haplotype is taken from a normal distribution centered around this average value. Then, each haplotype is assigned a certain sequence divergence, with average defined by the --divergence parameter. As for the haplotype size, also the haplotype divergence is taken from a normal distribution centered around the declared average sequence divergence. Using normal distributions to generate haplotype length and divergence ensures that most haplotypes will have a length and a divergence compatible with the ones requested by the user, but also ensures the presence of some outliers (e.g. a hyper-variable region) that help mimicking a real-life scenario. In a third step, the “sim” module turns some of the haplotypes into LOH blocks. This is done by turning the block’s sequence divergence to zero. The fraction of haplotypes that are turned into LOH blocks is defined by the --loh parameter. Subsequently, “jloh sim” adjusts the sequence divergence of the other haplotypes to bring back the average to the value declared --divergence parameter. This is important, because if the user wants 40% LOH, then the resulting sequence divergence will be lower than the one requested via the --divergence parameter. Once all these steps are completed, ”jloh sim” iterates through the haplotypes and introduces as many SNPs in random positions as are required to reach the sequence divergence assigned to each haplotype. The haplotypes, together with their connotation (“normal” or “LOH”) are reported in an output file alongside the FASTA file representing the divergent genome. Note that this process generates only heterozygous SNPs.

This module was run 24 times, each with a different combination of sequence divergence and LOH rate as mentioned above. Then, each of these 24 diverging genome versions was concatenated with the original *S. cerevisiae* S288C genome sequence and reads were simulated from the concatenated version. These simulated reads represented simulated hybrid reads. A number of reads was estimated in order to reach approximately 10X and 30X coverage of a *S. cerevisiae* hybrid genome. The hybrid was assumed to have twice the genome size of *S. cerevisiae*, i.e. 12 Mbp x 2 = 24 Mbp. The number of paired-end reads of length 125 bp (i.e. 2×125=250 bp) required to reach ∼10X and ∼30X coverage over a 24 Mbp genome size was calculated, returning approximately 1 million and 3 million reads. These values were then used to simulate reads in wgsim (https://github.com/lh3/wgsim) as the “-N” parameter, with the following other parameters: -d 1000 -s 200 −1 125 −2 125. Combining the divergence rate, the LOH rate, and the simulated coverage, the simulated dataset was composed of 48 simulated samples.

Long PacBio RS-II reads were simulated with pgsim v3.0.0 (27) using mostly default parameters except --strategy wgs --method errhmm --errhmm data/ERRHMM-RSII.model --depth 30.0. This set of parameters returned simulated PacBio reads at 30X coverage. Reads were simulated from the simulated genomes carrying 15% and 20% divergence with the *S. cerevisiae* genome sequence (see above).

### 2.5 Testing read mapping impact

The SNPs that “jloh extract” uses to infer LOH blocks are derived from short read mapping. When assessing the degree of LOH, the sequence divergence between reads and reference genome is a key factor. Hence, the limits of read mapping were tested in relationship to the sequence divergence. A genome with 20% sequence divergence from the *S. cerevisiae* genome sequence was simulated using “jloh sim”. It then was concatenated to the original *S. cerevisiae* genome and 125 nt paired-end reads were simulated from this concatenation to imitate a genome with two very divergent sets of haplotypes (e.g. a hybrid) using the same simulation parameters described above. Reads were then mapped onto the *S. cerevisiae* genome alone, using two common mapping software: bowtie2 and BLAT (28, 29). Reads were mapped with both tools with progressively less stringent minimum sequence identity between reads and reference (from 100% to 70% in steps of 5%). In the case of bowtie2 this is controlled by the --score-min and --mp parameters, which were set to --score-min L,0.0,-x and --mp 6,2, where x is a fraction of the read length. To achieve the required sequence identity thresholds, the “x” in --score-min was set to 0.0, 0.3, 0.6, 0.9, 1.2, 1.5, and 1.8. In the case of BLAT, the -minIdentity parameter allowed a more direct control over the minimum sequence identity. The amount of reads that were mapped in each run by each program was then quantified and used as measurement for read mapping reliability under the assumption that the perfect scenario is where all reads get to be mapped and heterozygous SNPs emerge from divergent regions.

### 2.6 Read mapping and variant calling

Simulated short reads were mapped against the reference genomes of their parental species independently. The mapping was conducted with HISAT2 v2.1.0 (30) using very relaxed parameters (--score-min L,0.0,-1.0 --mp 6,2 --rfg 5,3 --rdg 5,3 -I 0 -X 1000). Simulated long reads were mapped with Minimap2 (31) with parameters -H -x map-pb -a. The following steps of the workflow were applied to both short and long reads. The mapping records were filtered and sorted by genome coordinate using samtools v1.15 (32), removing secondary alignments and unmapped reads (-F 0×0100 -F 0×4). Filtered mapping records were used to call short variants using bcftools v1.15 (33). First, reads were piled up using bcftools pileup (--annotate FORMAT/AD,FORMAT/ADF,FORMAT/ADR,FORMAT/DP,FORMAT/SP,INFO/AD,INFO/ADF,I NFO/ADR --output-type v --skip-indels). Then, pileups were used to perform SNP calling using bcftools call (--multiallelic-caller –variants-only); indels were not considered. Raw SNPs were filtered removing those with QUAL ≥ 20, AF ≥ 0.05, DP ≥ 4, and MQ0F ≤ 0.05. This filtering was conducted with all2vcf (https://github.com/MatteoSchiavinato/all2vcf). The final lists of SNPs from either parent of a hybrid were passed to JLOH to detect LOH blocks.

### 2.7 LOH block inference sensitivity and specificity

To test the sensitivity and specificity of “jloh extract”, a series of runs were performed with each of the 48 simulated data sets described above. Some parameters were kept fixed in all runs: SNPs were considered heterozygous if their alternative allele frequency (AF) was within 0.3 an 0.7 (--min-af 0.3 --max-af 0.7); candidate blocks were required to be at least 50% covered by reads (--min-frac-cov 0.5); blocks were considered hemizygous if their average coverage per position was lower or equal to 75% of their 5000 bp flanking regions (--hemi 0.75 --overhang 5000). The variable parameters were instead the minimum block length (--min-length) and the SNP density thresholds (--min-snps-kbp). The --min-length parameter was set at 500, 1000, 2000, 5000 or 10000 bp. The SNP density thresholds were set to the 5th, 10th, 15th and 50th quantile as detected via “jloh stats” (see above) A total of 960 runs represented all possible combinations between simulated samples and parameters. From each run, the true positives (TP), true negatives (TN), false positives (FP) and false negatives (FN) were assessed. We defined them as follows. TP: genome positions that are part of an LOH block introduced upon simulation, and are also within a block found by “jloh extract”. FP: genome positions that are not part of a simulated LOH block, but are within a block found by “jloh extract”. TN: genome positions that are not part of a simulated LOH block, and are also not within a block found by “jloh extract”. FN: genome positions that are part of an LOH block introduced upon simulation, but are not within a block found by ”jloh extract”. Consequently, the sensitivity or true positive rate (TP rate) is obtained via TP / (TP + FN) and the specificity or true negative rate (TN rate) is obtained via TN / (TN + FP). These values oscillate between 0 and 1. Runs with both metrics above 0.75 were considered “good”, while runs with both metrics above 0.90 were considered “excellent”. All these calculations were done with the custom python script “tp_tn_rates.py” available within the JLOH github repository (https://github.com/Gabaldonlab/jloh).

### 2.8 Reanalysis of public data

To further test the potential of the “jloh extract” module, data from public sequencing libraries compiled in a previous study (18) was downloaded. Among these samples, only five had their LOH blocks called (22). These five samples are available in SRA under the SRA codes ERR1111502, ERR1111503, ERR3010130, SRR2967902, and SRR2967903. All these samples represent hybrids between *S. cerevisiae* and *S. paradoxus*. To better identify the strains in the results their strain name was used. Respectively: UFMGCMY651, UFMGCMY652, CBS7002, AWRI1501 and AWRI1502. Reads from these five samples were preprocessed using Trimmomatic (v0.39, Bolger et al., 2014). Trimming was conducted using the following parameters on all the read files: LEADING:20 TRAILING:20 SLIDINGWINDOW:4:25 AVGQUAL:20 MINLEN:35. Accession codes and related genomic assignments can be found in Supplementary File S1. Reference genomes and versions used here were the same used by Bendixsen et al. (2021).

To map reads and extract SNPs the same workflow described above for simulated data was used. The same *S. cerevisiae* reference genome sequence described above was used in conjunction with the ASM207905v1 assembly of the CBS432 strain of *S. paradoxus* (35). The generated VCFs and BAMs were passed to “jloh extract” to infer LOH blocks with the same fixed parameters described for the simulated data. In addition, we set the following two parameters: --min-length 1000 --min-snps-kbp 15,3.

The mapped reads in BAM format were used to extract read coverage per position over all chromosomes of the *S. cerevisiae* and the *S. paradoxus* parental genomes. The coverage information was then binned in windows of 10,000 bp, calculating the mean coverage per position of each window. The difference in coverage between the two groups of windows was assessed using a Mann-Whitney test (signif. p < 0.05, two-sided). The LOH blocks inferred with “jloh extract” on these two parental genomes with the same reads and the derived SNPs were then crossed with these 10 kbp windows. We attempted to assign each window to either *S. cerevisiae* or *S. paradoxus* depending on the amount of basepairs in its span that overlapped LOH blocks from either one of the two parents. The assignment of windows along each chromosome was then plotted using a custom R script. Genes that could be involved in the LOH anomaly were extracted using *S. cerevisiae* transcript sequences as template. These sequences were mapped against the *S. paradoxus* genome sequence using gmap (-f gff3_gene) (36). Transcripts mapping in positions 1 to 370,000 of the *S. paradoxus* chromosome 12 were extracted, and their corresponding protein sequences were submitted to the Pannzer2 webserver for GO annotation (37). The list of gene symbols and relative GO terms was analysed in R using clusterProfiler (38). A list of EntrezIDs was obtained from the gene symbols, and the enrichment of their corresponding GO terms was assessed with the enrichGO() function (ont = “ALL”, pvalueCutoff = 0.05, pAdjustMethod = “BH”, qvalueCutoff = 0.2, minGSSize = 5, maxGSSize = 500).

### 2.9 LOH block inference in *C. orthopsilosis* clades

To show the potential of “jloh extract” in detecting differences in LOH blocks from different clades of the same hybrid species, a set of sequencing data from the hybrid fungal pathogen *C. orthopsilosis* was used. The strains represented in the data belonged to four separate clades of *C. orthopsilosis*. Specifically, the usefulness of “jloh stats” and SNP density quantiles in distinguishing groups of samples by inferred LOH was assessed. Sequencing libraries used were retrieved from BioProjects PRJEB4430 and PRJNA322245 (39, 40). The ASM31587v1 version (41) of the *C. orthopsilosis* reference genome sequence was used for the read mapping and LOH block inference. Reads were trimmed using Trimmomatic v0.39 (34). Mapping and variant calling was performed using PerSVade v.1.02.4 using 5000 as minimum chromosome length and a value of 20 as minimum coverage. Variants passing quality filters by at least two out of the tree software that PerSVade uses were included in the analysis.

The “jloh stats” module was run using genomic variants extracted from strains belonging to each of the four clades of *C. orthopsilosis* hybrids. The SNPs/kbp quantiles were collected for all strains, and an average quantile value per clade was calculated for quantiles Q5, Q10, Q15, Q50, Q85, Q90 and Q95 (Supplementary File S2). The “jloh extract” module was then run to call LOH blocks for all samples using the listed quantiles to set the --min-snps-kbp parameter. Candidate blocks were required to be 90% covered by reads (--min-frac-cov 0.9) and only SNPs annotated as “PASS” were kept (--filter-mode pass). Bedtools v2.30.0 multiinter (42) was then used to find overlapping LOH blocks between all strains and between strains belonging to the same clade. Files harbouring LOH blocks were loaded and visually examined with IGV v2.8.13 (43).

### 2.10 LOH block inference in *C. metapsilosis*

Sequencing libraries for samples BP57 (SZMC8022) and PL448 were retrieved from BioProject PRJNA238968 (17). The chimeric assembly of hybrid BP57 was used as reference genome (Mixão *et al.*, *under revision*). Read filtering, mapping and variant calling was performed as described in the section above. The “jloh stats” module was run prior to LOH block inference in order to obtain the optimal SNP density parameters for each of the two strains. The “jloh extract” module was then run using Q50 quantile values in the --min-snps-kbp parameter. Candidate blocks were required to be 90% covered by reads (--min-frac-cov 0.9) and only SNPs annotated as “PASS” were kept (--filter-mode pass). The “jloh plot” module was used to visualise the resulting LOH blocks.

### 2.11 Code specifics

The core algorithm of JLOH is coded in python (v3.9.7) using the pybedtools v0.8.2 module for most interval-related operations (44) and self-written python functions for deeper analyses. The analysis carried on in this study has been performed with a pipeline written in nextflow v21.04.3 (45) which went from read mapping to LOH block extraction. The pipeline containing the data simulation and fidelity testing steps, together with its configuration file and a shell script to run it, is provided in the “workflows” directory of JLOH’s github repository (https://github.com/Gabaldonlab/jloh). The pipeline has been run on a single node of a computing cluster. The node had 48 processors (2.1 GHz each) and 94 GB RAM.

## 3. Results

### 3.1 Read mapping strongly influences results fidelity

Mapping is most often conducted with default parameters using tools such as bowtie2, hisat2, bwa or BLAT (28, 29, 46, 47). The first three are common fast aligners that use a seed-and-extend strategy to map reads. In order to be precise and reliable, seeds must not be shorter than 18-20 bp. The latter, instead, is a command-line tool equipped with the same algorithm of BLAST and which searches for matches of any small tag (8-12 bp) from the query sequences (the reads) into the provided reference genome, extending each match and providing a score. These programs have been written and optimized to map reads from one species against the genome of the same species, but when the sequence identity between reads and reference is beyond the one expected in an intra-specific scenario, they may fail to properly map all reads in the dataset. This is due to inherent limits within their algorithms, which are based on exact seed matches that cannot have mismatches with the reference genome. In the presence of reads that are highly divergent from the reference genome, an exact seed to extend the alignment may not be found. For example, if the reads are on average 20% divergent from the reference genome sequence, there will be an average of 20 SNPs per 100 bp, which in a seed of 20 bp turns into 4 mismatches. To demonstrate this limit we used the genome of *S. cerevisiae* to simulate a diploid set of reads with an internal 20% divergence between its two sets of chromosomes (see methods). We conducted the simulation with “jloh sim”, which breaks the genome sequence into a random assortment of haplotypes each with its own divergence against the reference genome sequence (see methods). Hence, while the average divergence of the simulated dataset was 20%, it also contained haplotypes with lower or greater divergences. We then mapped this read set back onto the *S. cerevisiae* genome using bowtie2 and BLAT. With both tools we performed several runs at progressively more relaxed criteria in terms of minimum accepted sequence identity between reads and reference (from 100% to 70% in steps of 5%); results are shown in Table 1.

**Table 1.**
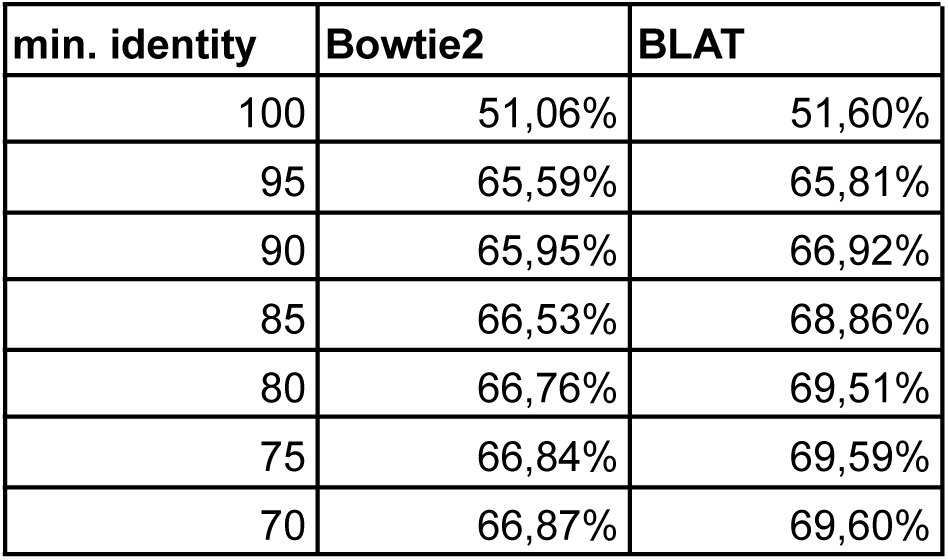
Mapping rates obtained with either Bowtie2 or BLAT at different thresholds of minimum sequence identity.

| min. identity | Bowtie2 | BLAT |
| --- | --- | --- |
| 100 | 51,06% | 51,60% |
| 95 | 65,59% | 65,81% |
| 90 | 65,95% | 66,92% |
| 85 | 66,53% | 68,86% |
| 80 | 66,76% | 69,51% |
| 75 | 66,84% | 69,59% |
| 70 | 66,87% | 69,60% |

As expected, when allowing exact matches only (i.e. 100% identity), both bowtie2 and BLAT managed to map only half of the reads against the *S. cerevisiae* genome (51.06% and 51.6% mapping rate, respectively). When relaxing the parameters down to 70% minimum identity between reads and reference, which would theoretically be sufficient to capture 20% divergence between subgenomes, both software still failed to map more than 30% of the reads. A quick look at the coverage per position showed that the haplotypes with the least coverage were also having the highest divergence between *S. cerevisiae* and the simulated copy used to generate the reads. Hence, we concluded that read mappers fail to identify good alignment seeds to extend reads in these regions, therefore leaving them uncovered. We stress that this is important when working with LOH blocks as well as anything that depends on read mapping, since real-world genomes may have hypermutated regions that may not be captured with default mapping parameters.

### 3.2 Sensitivity and specificity

We benchmarked the accuracy of “jloh extract” in detecting LOH blocks from Illumina sequencing data. We set up a test that included the simulation of data containing known LOH blocks and the retrieval of LOH blocks with “jloh extract” combined with “jloh stats” to study the SNPs/kbp distribution. We first used “jloh sim” to simulate a series of yeast hybrid genomes carrying different levels of allelic divergence between homeologous chromosomes (1%, 3%, 5%, 10%, 15%, 20%) and different amounts of LOH (10%, 20%, 30%, 40% of the genome). The LOH blocks introduced in the simulation defined the true positives of the test, as their location was known. Then, from each of these genomes we simulated short sequencing reads, mapped them against the *S. cerevisiae* genome sequence, and called SNPs. We passed the called SNPs to “jloh stats” to detect the chosen SNP distribution quantiles (Q5, Q10, Q15, Q50), and we then ran “jloh extract” multiple times (one per relevant quantile) to retrieve LOH blocks. The inferred LOH blocks were then compared with the known simulated blocks. We note that the fact that “sim” and “extract” are both part of the same toolkit is merely for practical reasons but they are independent programs that do not have any intercommunication and therefore cannot influence each other’s results. To assess the impact of different parameter choices, we repeated this benchmark over different parameter settings. The tested parameters were: 1) the minimum length to retain a candidate LOH block (500, 1000, 2000, 5000 and 10000 bp); 2) the simulated sequencing depth (10X or 30X); 3) the SNP density quantile to use as threshold to isolate candidate LOH blocks (Q5, Q10, Q15, Q50). We note that the last parameter is a combination of the average heterozygous and homozygous SNP densities, it is generated by “jloh stats”, and is passed to the --min-snps-kbp parameter of “jloh extract”. All possible combinations of these parameters, the LOH rates, and the divergence rates resulted in 960 runs.

We calculated the true positive rate (TP rate, or sensitivity) and the true negative rate (TN rate, or specificity) from each of the runs (Supplementary Figure S1, Supplementary File S3). The sensitivity reflects the ability of JLOH to identify *bona fide* LOH blocks (true positives), while the specificity reflects its ability to avoid reporting false positives in the results. All 960 runs had sensitivity ≥ 0.90, of which a notable 93.3% reached a sensitivity of 0.99, attesting to the ability of “jloh extract” to find the LOH blocks that were introduced from the simulation script. When looking at specificity, which is higher in absence of false positives, 68.3% of the run had it ≥ 0.90. This means that, combining sensitivity and specificity, 68.3% of the runs had both statistics ≥ 0.90 (Supplementary Figure S1).

To investigate further why certain runs had low specificity we categorized JLOH’s performance in each run by considering as “good” the runs that had both sensitivity and specificity ≥ 0.75, and as “excellent” those that had both ≥ 0.90 (Supplementary File S4). A quick look at the run specifics showed us that the runs with the most false positives (i.e. those affecting specificity) were those where the simulated divergence was higher (15% or 20%), while other parameters did not particularly influence the quality of the results (Figure 3, Supplementary Figure S2, S3, S4, S5).

**Figure 3.**
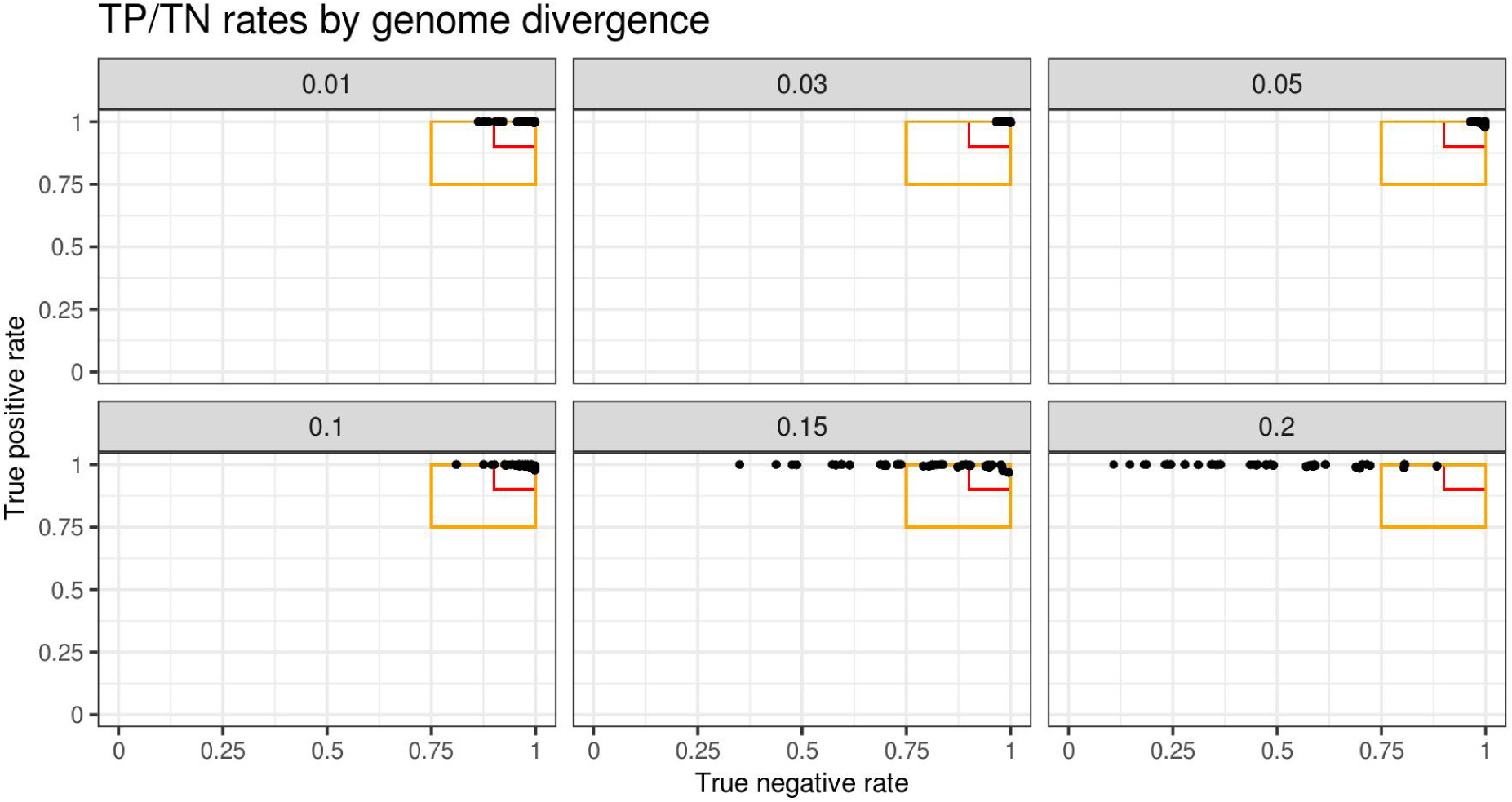
Sensitivity (TP) vs specificity (TN) rates in all the runs performed with simulated data, subdivided in facets corresponding to the simulated sequence divergence between subgenomes of the simulated hybrid data. Divergence rates are indicated from 0.01 (1%) to 0.20 (20%). The orange box in each facet represents “good” runs (TP and TN ≥ 0.75) while the red box represents “excellent” runs (TP and TN ≥ 0.9).

At 1% divergence 90% of the runs were excellent and 100% were good. At 3% and 5% divergence all runs were excellent. At 10% divergence, 92.5% of the runs were excellent, meaning that “jloh extract” is very capable of inferring LOH blocks in the 1-10% divergence range, and the results carry little false positives. At 15% divergence, however, only 27.5% of the runs were excellent. Of these 44 excellent runs, 36 were performed with very long minimum block size (5000-10000 bp), and 28 with higher (30X) simulated sequencing depth. Also, the algorithm’s precision lowered linearly with the increasing of the simulated LOH rate in the divergent genome, with 24 out of 44 runs being from 10% LOH rate, 12 from 20%, 8 from 30%, and none from 40%. This suggests that increasing divergences lead to patches of the genome left uncovered due to the read mapping issues described in the previous section, which emerge especially with short reads. Nearly all the introduced LOH blocks are recovered even at high divergence, but many false positives were introduced which lower the specificity score. The minimum length criterion of selection partially quenches this effect by weeding out short uncovered patches, but the results become less reliable. These effects were amplified when looking at runs performed with reads derived from genomes that were 20% divergent. None of the runs were excellent, and only 10% were good. Of these good runs, almost all of them were obtained from 30X simulated depth reads, 10000 bp minimum block length, and low LOH rates (10% or 20%). All in all, we concluded that there is a technical limit to reliable LOH block inference based on short read mapping, and this limit is placed around 15% sequence divergence, above which a fraction of reads will fail to find a seed to extend due to excessive divergence.

To prove that read length is responsible for this effect we simulated PacBio RSII long reads from the 15% and 20% divergent simulated genomes described above combined with the *S. cerevisiae* genome sequence. This combination mimics the context of a 15% or 20% divergent hybrid, as described above for the short reads experiment. These two genomes carry LOH blocks introduced by the “jloh sim” module of our program, hence allowing us to estimate true positive and true negative rates. We mapped these simulated hybrid reads against the *S. cerevisiae* genome sequence, called SNPs and used them to infer LOH blocks with “jloh extract”. Indeed, the true positive rate and true negative rate increased substantially. The 15% divergent run returned a TP rate of 1 and a TN rate of 0.93. The 20% run, instead, returned a TP rate of 1 and a TN rate of 0.91. In both cases, the results obtained were substantially better than the ones obtained with short reads. Hence, we concluded that using long reads solves the mapping issues that hamper LOH inference in hybrids from highly divergent parents.

We then attempted to compare JLOH’s efficacy in retrieving LOH blocks against existing software, namely MuloYDH (22) and YMAP (21). However, YMAP website was unavailable (connection time-out error, last accessed 31 March 2023), and MuloYDH failed to complete the run due to multiple errors or returned empty output files. Hence, a comparison with existing software was unfortunately not possible, which underscores the need for a general purpose solution. All in all, based on our benchmark, we concluded that nearly 70% of the parameter and testing space resulted in very reliable LOH block detection, and that sequence divergence between reads and reference is the most influential factor in affecting the fidelity of results.

### 3.3 Scalability

We assessed the scalability of JLOH in terms of CPU usage, and the impact of the dataset size on the computational time. The latter refers to the ability of JLOH to tackle progressively larger sets of SNPs without losing efficiency. To perform this test, we generated a simulated, divergent copy of the *S. cerevisiae* genome using the “sim” module of JLOH. The copy had 10% sequence divergence and 30% of its sequence was carrying simulated LOH blocks, with a mean simulated block length of 5000 bp. Then, we simulated reads from these two genomes following the same procedure described in previous paragraphs, and called SNPs with these reads by mapping them against the *S. cerevisiae* genome and the divergent simulated copy of it. These SNPs, their source SAM file, and the reference genomes were the inputs for our scalability testing. The dataset was composed of 697,494 SNPs inferred from mapping 3 * 10^6^ read pairs of length 2 x 125 nt over the combination of the *S. cerevisiae* genome and the simulated divergent copy (24 Mbp in total).

To test the scalability in terms of CPU usage, we ran the “extract” module with the whole dataset as input in separate jobs using from 1 to 48 cores, in steps of 4. Results are shown in Supplementary Figure S6, panel “A”.

To test the dataset size impact on computational time, we formed four subsets from the previously mentioned simulated data. These subsets represented: 1) the whole genome composed of 16 chromosomes + the mitochondrial chromosome; 2) chromosomes I-IV; 3) chromosomes I-VIII; 4) chromosomes I-XII. Then, we ran the same JLOH extract command on each subset, and logged the elapsed computational time. Results are shown in Supplementary Figure S6, panel “B”. Results show that the computational time gain derived from CPU parallelization reaches a plateau at about 10 CPUs, while the computational time increases linearly with input data size.

### 3.4 Estimating appropriate SNP density thresholds

As we discussed previously, the parameter regulating the minimum density of heterozygous and homozygous SNPs (--min-snps-kbp) for candidate blocks is key for a successful block inference. The “jloh stats” module aids the user in choosing the right setting for this parameter by modelling the SNP density distribution and providing quantiles that the user may use as thresholds. To test the robustness of this idea we used hybrids from *C. orthopsilosis*, a species of emergent fungal pathogens that belong to the *C. parapsilosis* species complex (48). The majority of isolates sequenced to date are hybrids with the exception of one homozygous (parental) lineage (39). The second parent remains unknown. All *C. orthopsilosis* hybrids descend from four independent hybridisation events between the same two parents which are approximately 5% divergent (Figure 4) (39, 40).

**Figure 4.**
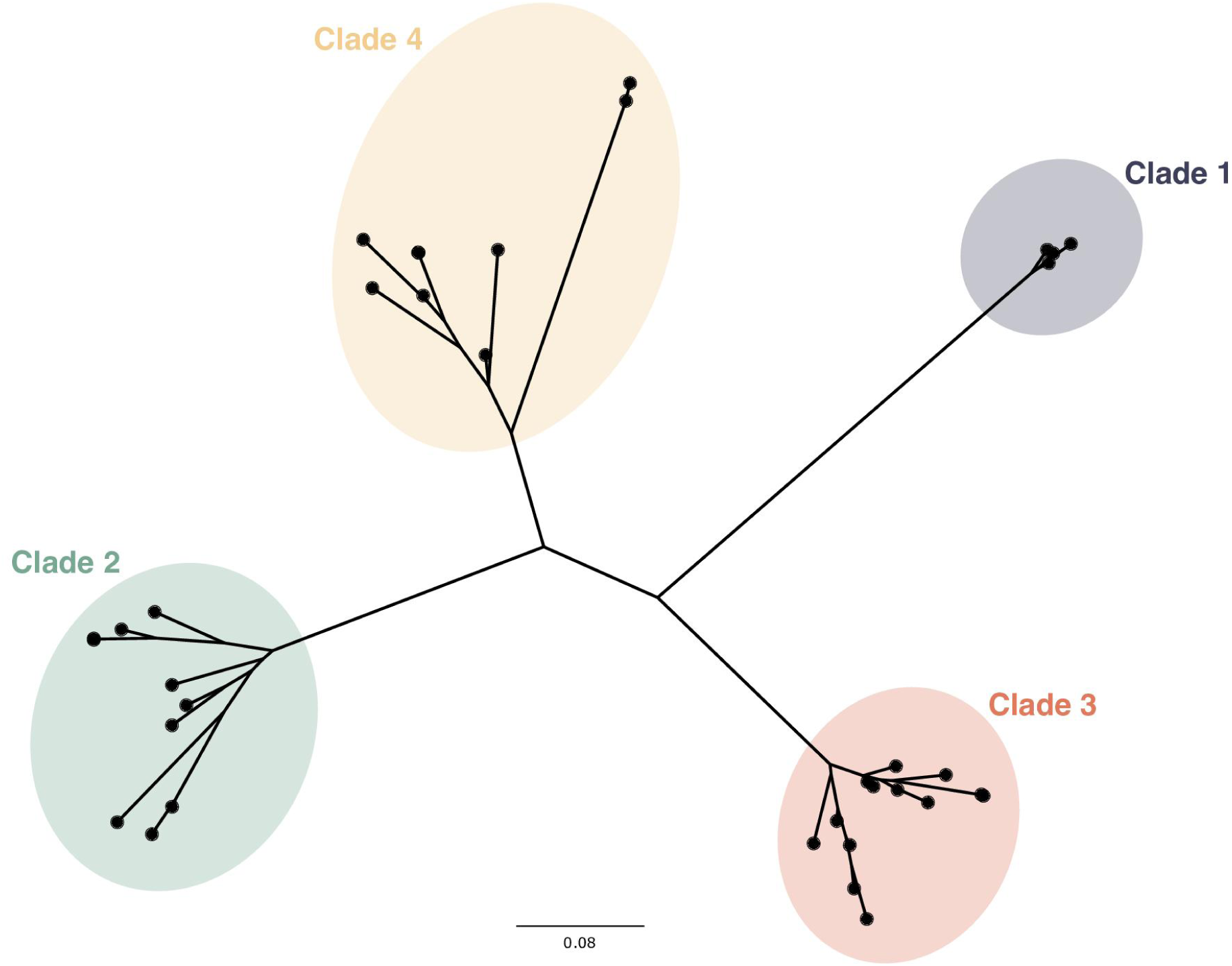
Tree based on variants depicting phylogenetic relationships between *C. orthopsilosis* hybrid strains. The four known clades are marked.

The four hybridisation events gave rise to four hybrid clades which are very different in terms of SNP density and patterns of LOH (40). These differences make *C. orthopsilosis* hybrids a good case study to test how setting different minimum SNPs/kbp thresholds may influence the efficiency of LOH block calling.

In general, average block size was inversely proportional to the quantile value and Q50 yielded the highest amount of blocks for all hybrids (Figure 5, panel “A”; Figure 4; Supplementary File S2; Supplementary Figure S7).

**Figure 5.**
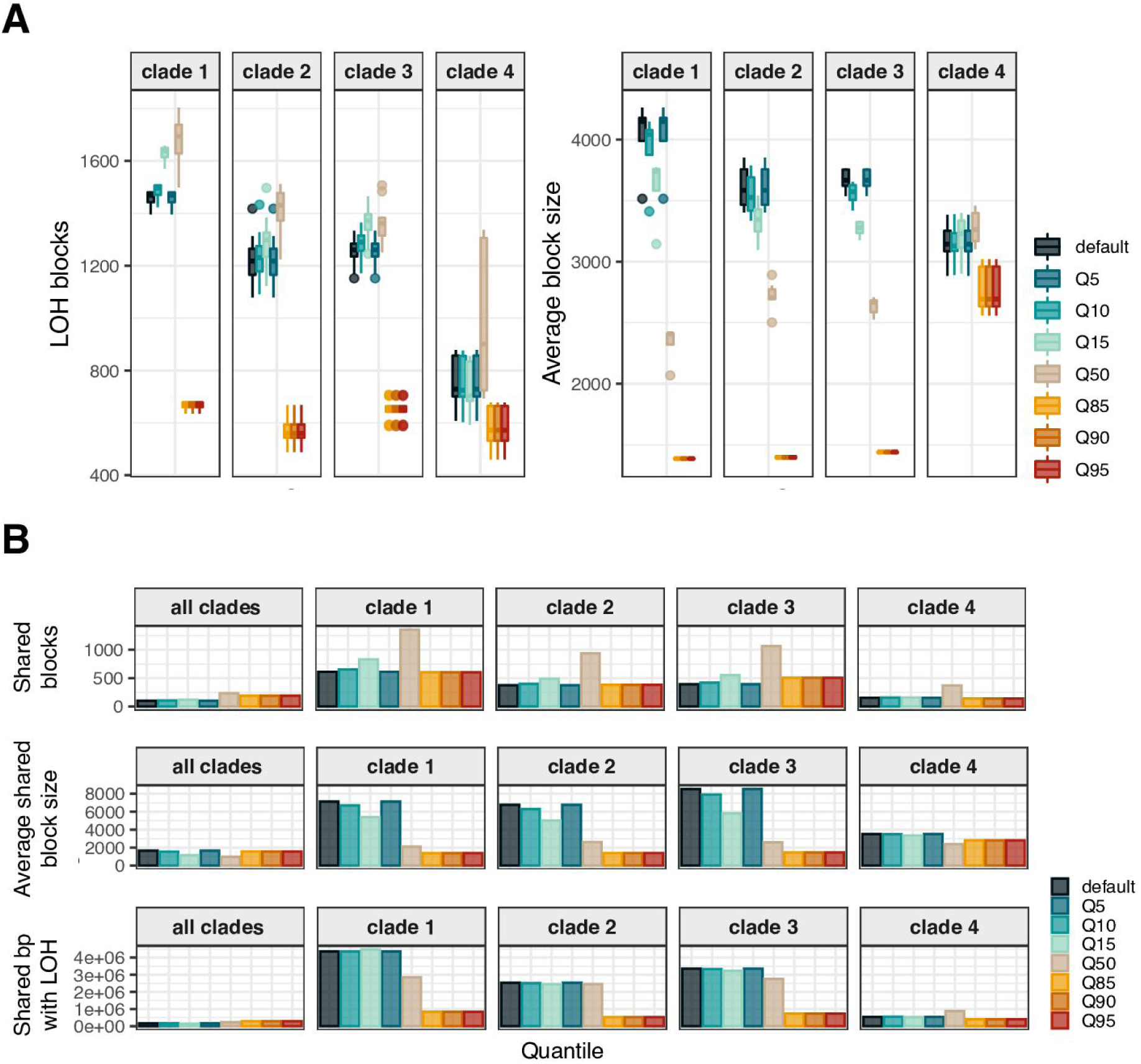
The number of inferred LOH blocks and their size depend on the different quantile values used (A) number and size of LOH blocks in all *C. orthopsilosis* hybrid strains using different quantile values. (B) Number, average size and amount of base pairs of shared LOH between four representative strains of each clade.

Each of the four different hybridisation events between *C. orthopsilosis* parental strains gave rise to a different clade of hybrids. Because of this shared evolutionary history, hybrids that belong to the same clade share similar LOH patterns (i.e. specific coordinates of LOH blocks), which differ from those of hybrids belonging to another clade. We took advantage of this clade specificity to assess which parameters would be optimal for LOH inference, as a good parameter set is expected to reflect the clade of origin for each strain whereas false positives and negatives would blur the clade-specific evolutionary signal. We selected four representative strains of each clade and assessed the overlap between LOH blocks called within and between clades. Parameter values leading to the highest overlap of blocks between samples of the same clade without increasing the overlap between different clades were considered most suitable (Figure 5, panel “B”, first row). Our results clearly show that blocks called with parameters set at the Q50 quantile of SNPs/kbp resulted in the highest specificity in detecting shared LOH blocks between strains of the same clade (Figure 5, panel “B”; Supplementary File S5).

In sum, the number of minimum SNPs per kilobase considered to define blocks does influence the results of the LOH block inference. Therefore, the “jloh stats” module represents a helpful tool to optimise LOH block calling parameters according to specific experimental settings and scientific question, such as the quest for shared blocks between strains that we used in this paragraph.

### 3.5 Reanalysing public variation data

With the provided framework, it is possible to extract LOH blocks from pre-existing variation SNP data that was used for other purposes. To demonstrate the advantage of JLOH in this context, we processed five genome sequencing samples from a dataset of publicly available hybrid yeasts (18) taken from a collection of 204 publicly available yeast hybrid samples which the authors characterized in terms of their environment (e.g. beer, wine, olive). Within that study, the authors extract LOH from five *S. cerevisiae* x *S. paradoxus* hybrids only. These hybrids were generated in three separate original studies (49–51). These strains were UFMGCMY651, UFMGCMY652, CBS7002, AWRI1501 and AWRI1502. By making a consensus of the LOH blocks called in the five hybrids, the authors extract an LOH propensity for each region of each chromosome, and show that the majority of chromosome 12 was showing a remarkably strong signal of LOH favouring the *S. cerevisiae* alleles (Figure 8 in the original publication). Our results confirm this trend but also show further information when stratified by individual strains. When looking at the LOH propensity of chromosome 12 of *S. cerevisiae* (Figure 6, panel “A”), four out of the five strains show the expected *S. cerevisiae* predominance in terms of haplotype, but one (CBS7002) shows an LOH profile that is almost entirely favouring *S. paradoxus* alleles. This trend would not be visible when making a consensus of the five strains, as the other four have a strong coherent signal that would skew the consensus result towards *S. cerevisiae*’s alleles. We then looked at the complementary results that we obtained on chromosome 12 of *S. paradoxus* (Figure 6, panel “B”). These results look coherent with the ones obtained on *S. cerevisiae*, as the same four strains show LOH predominantly favouring *S. cerevisiae* alleles. Strain CBS7002, as in Figure 6A, displays a different trend.

**Figure 6.**
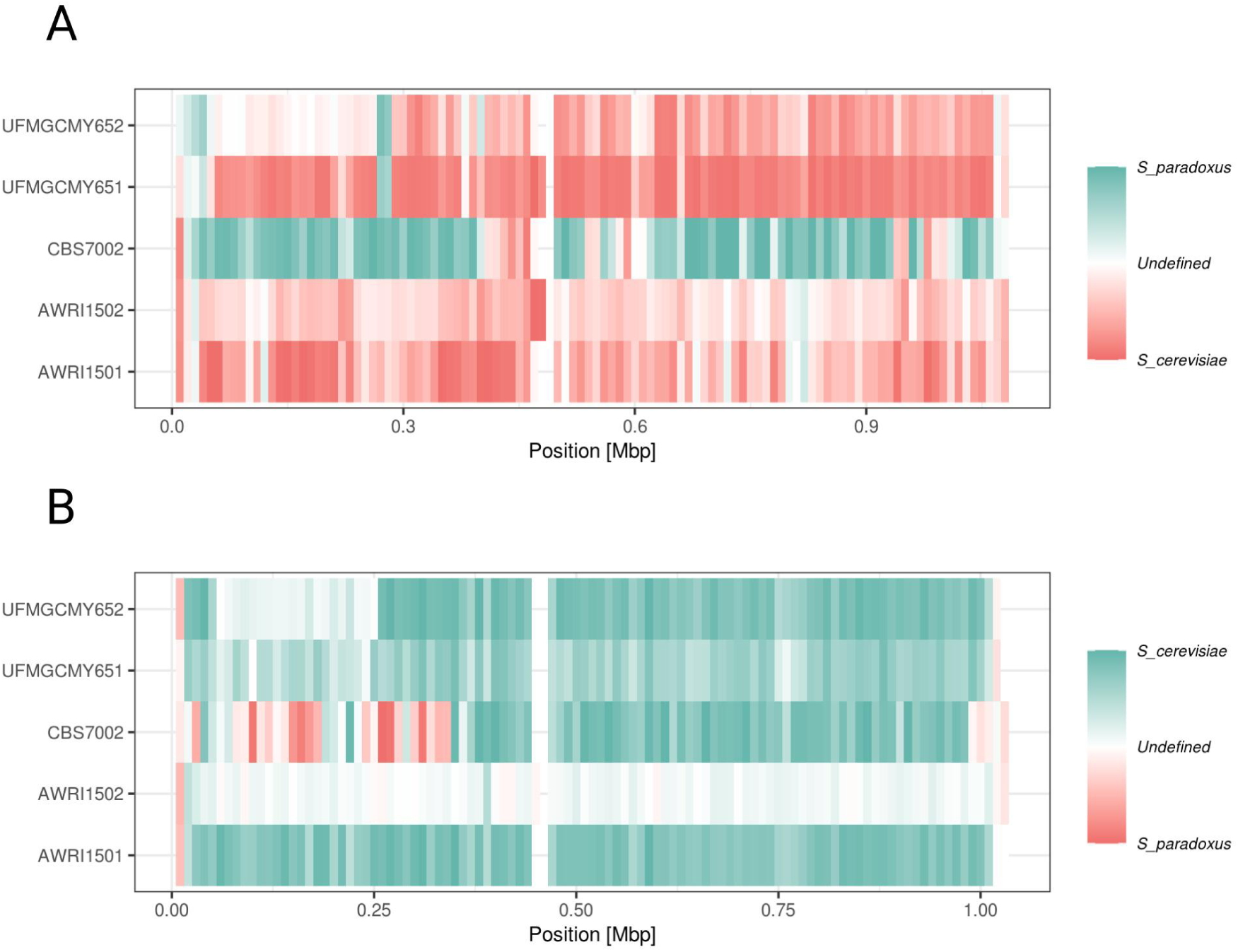
Density of LOH signal along chromosome 12 of *S. cerevisiae* (**A**) and of *S. paradoxus* (**B**) computed in windows of 10 kb in five *S. cerevisiae x S. paradoxus* hybrids. Colors represent allele assignment (REF or ALT) while color intensity represents the fraction of positions in each window that were in a predicted LOH block. Note that in (**A**) “blue” corresponds to *S. paradoxus* alleles while in (**B**) it corresponds to *S. cerevisiae* alleles, as it generally represents the alternative allele.

The initial part of chromosome 12 (approx. 1-370 kbp) shows LOH favouring *S. paradoxus* in line with the results observed on chromosome 12 of *S. cerevisiae*, while the remaining part of the chromosome shows LOH towards *S. cerevisiae*. This is, of course, impossible: in both cases the suggested LOH allele is the alternative allele (either *S. cerevisiae* or *S. paradoxus* depending on the chromosome), which indicates the possibility that a third genotype is in play in this scenario, which is neither the *S. cerevisiae* nor the *S. paradoxus* genome used for the analysis. Sample CBS7002 comes from a study where researchers studied a quasi-domesticated hybrid strain of *S. cerevisiae x S. paradoxus* adapted to an olive industrial processing environment (51). The authors describe it as a hybrid which lost most of its *S. paradoxus* subgenome due to LOH in favour of the *S. cerevisiae* subgenome. Figure 6 clearly shows how, in this isolate, the first portion of chromosome 12 has instead fully retained the *S. paradoxus* alleles. We speculate that the alleles carried by the *S. paradoxus* subgenome in this region of chromosome 12 may have been more advantageous for this particular strain and were preferentially retained. We analysed the gene content of the first 370,000 positions of chromosome 12. Associated GO terms showed a high abundance of transmembrane proteins involved in membrane transport, although no GO term was significantly enriched. These types of proteins have been described as enriched in this hybrid by the original authors (51).

Another interesting deduction that we make from this analysis regards samples AWRI1501 and AWRI1502, which show diverging LOH patterns despite coming from the same study (50). Both AWRI1501 and AWRI1502 samples are hybrid yeasts commercialized from the Australian company Maurivin (https://www.maurivin.com/). However, while it was used as a *S. cerevisiae x S. paradoxus* hybrid in Bendixsen et al. (2021), according to Maurivin AWRI1502 is actually a *S. cerevisiae x S. cariocanus* hybrid. *S. cariocanus* is phylogenetically close to *S. paradoxus* and has been recognized as an independent species about 20 years ago (52). This attests to the ability of JLOH to distinguish LOH blocks even down to individual strains, and all in all, we concluded that JLOH is able to infer reliable LOH blocks and aid in generating biologically relevant knowledge.

## 4. Discussion

In this study we introduce JLOH, a computational toolkitl for inferring and analysing LOH blocks from SNP calling and read mapping data. To the best of our knowledge, only two tools exist that allow the extraction of LOH blocks starting from sequencing data, both focusing specifically on yeasts (21, 22). However, given their discussed limitations, a general-purpose, parallelizable, and straightforward command-line tool requiring common input such as FASTA, BAM and VCF formatted files was timely.

We here described the design and rationale of JLOH algorithms and showcase its applicability. We first demonstrated that the level of sequence divergence sensibly affected the quality of the retrieved LOH blocks. We stress that this divergence may be interpreted in two ways: on one hand it could represent the divergence between the sequenced read set and the reference genome in a normal, non-hybrid organism. On the other hand, if working with a hybrid species, it represents the divergence between the two subgenomes. At 1-10% divergence, compatible with intra-species diversity as well as the subgenome divergence of many known inter-species hybrids (12, 17, 39, 40, 53–56), JLOH was able to retrieve almost all the known LOH blocks with great precision. At higher divergence (15-20%) instead, we noticed a big impact of read mapping onto the results. In fact, we established a connection between the high number of true positives generated by “jloh extract” and the inability of read mappers to map all the reads when working with short reads. In situations where read mappers fail to map reads from one subgenome onto the other, “jloh extract” will infer a hemizygous block that lost one of the two alleles, as the read coverage halves in such areas. Hence, we established that when the reads and the reference genome have a sequence divergence of 15% or more, LOH blocks inferred from short read mapping become unreliable. This limit is determined by the read mapping algorithms and their need for a seed identical sequence of sufficient length. Through our simulation of PacBio RSII long reads, we demonstrated that longer reads alleviate the issue and improve LOH detection in hybrids with highly divergent sub-genomes. Future studies will have to take this into account when planning variant calling experiments on highly divergent genomes.

One of the research areas in which the study of LOH is of particular relevance is the analysis of hybrid genomes, as it has been demonstrated that LOH may efficiently drive their adaptation (7). JLOH is particularly useful in this context, since it can work with a paired dataset representing variant and mapping information from a hybrid’s subgenomes and reference sequences from their parental genomes. The mapping has to be conducted onto the two parental genomes independently, generating two variant sets and two read mapping record sets. As shown in our results, it is important to account for the expected sequence divergence between subgenomes as it may sensibly affect the read mapping step and therefore the LOH block inference. Through homozygous SNPs, when run in --assign-blocks mode “jloh extract” will also annotate to which parental each LOH block belongs to. To the best of our knowledge, this is a novel aspect.

In many cases, one of the two parental species of a hybrid is unknown, and so its genome is likely unavailable. This, however, does not impede the usage of JLOH as long as the other parental genome is available. As demonstrated here with the *C. orthopsilosis* hybrids, it is still possible to retrieve LOH blocks by mapping the reads from the hybrid against the genome sequence of one of the parents. Heterozygous SNPs will still be annotated and homozygous SNPs will display whether the block has been inherited from the parent used as reference (REF) or from another allele (ALT). This applies to contexts where both parents are missing, too. To showcase this, we inferred LOH blocks in two strains (BP57 and PL448) of the known hybrid *Candida metapsilosis* (17) using the BP57 chimeric genome assembly as reference. Using this approach, we could distinguish between regions of LOH and heterozygosity despite not being able to infer the original haplotype of each block. As expected, given that the two strains belong to the same clade, they displayed largely similar patterns and distribution of LOH blocks. Notably, in the original study (17), Pryszcz *et al*. reported that in strain PL448 one chromosome had undergone LOH through its entire sequence. Our results recapitulate this finding: we observe a complete lack of heterozygosity in scaffold 6 of strain PL448 (Supplementary Figure S8, panel “B”). On the contrary, this scaffold was highly heterozygous in BP57 (Supplementary Figure S8, panel “A”). Hence, we suggest that JLOH may be used also in situations whereby only the hybrid genome sequence is available, still providing biologically relevant information.

We used the *C. orthopsilosis* dataset also to showcase the effect of SNP density thresholds. *C. orthopsilosis* is a yeast hybrid with known intra-species population diversity, and our dataset comprised four clades originated from independent hybridization events of the same parental lineages. We used “jloh stats” to calculate the density (SNPs/kbp) distribution of hetero- and homozygous SNPs over a genome sequence in windows of fixed size. We ran it for each sample and then averaged the SNP density distributions by clade. As shown in the code implementation, “jloh stats” returns the informative quantiles of the SNP density distribution of both hetero- and homozygous SNPs. For Q5, Q10, Q15, Q50, Q85, Q90 and Q95 we averaged the quantile value among samples by clade. Then, we used this value in “jloh extract” through the --min-snps-kbp parameter, which we have shown to impact the results due to its interconnection to SNP density and divergence. Our results (Table QUANTILES, Figure 5) showed a high degree of variability depending on the quantile used. In particular, when using Q50, we were able to capture clade-specific blocks and see differences in the clades without losing too much signal. While we stress that this is not the optimal quantile for any LOH-based analysis with JLOH, we suggest the user to select a quantile that is compatible with the experimental design. For example, if the analysed dataset does not have high intraspecific diversity, lower quantiles will allow the recovery of the majority of LOH blocks.

Clade-specific signal such as the one showed with *C. orthopsilosis* hybrids can be easily visualized with “jloh plot”, as we have shown by re-inferring LOH on five hybrids from *S. cerevisiae* and *S. paradoxus*. Simply displaying the inferred LOH blocks from these five strains showed how different each can be, as was the case for strain CBS7002. Future studies may leverage this simple workflow to highlight differences between clades, strains, or even individuals, and may uncover unknown differences or mislabellings in datasets with an intuitive visualization.

## 5. Data availability

No new data was generated for this study. All the code and related files are available at the JLOH github repository (https://github.com/Gabaldonlab/jloh) together with a nextflow workflow to generate a simulated test dataset to assess the tool’s functionalities.

## Supporting information

Supplementary File description

## Acknowledgments

We thank Leszek Pryszcz and Veronica Mixão for their original scripts, which have been the groundwork of this software. We thank Devin Bendixsen for kindly providing the results of their analysis published in 2021 to facilitate comparison with our results.

## 8. Author contributions

TG and MS designed the experiments. MS, TG and VdO modelled the computational problem. MS coded the software. MS and VNM tested the software. MS and VdO performed the analysis. TG supervised the project. All authors have read and accepted the manuscript.

