## Supplementary Material for "JLOH: Inferring Loss of Heterozygosity Blocks from Sequencing Data"

### Supplementary Figures

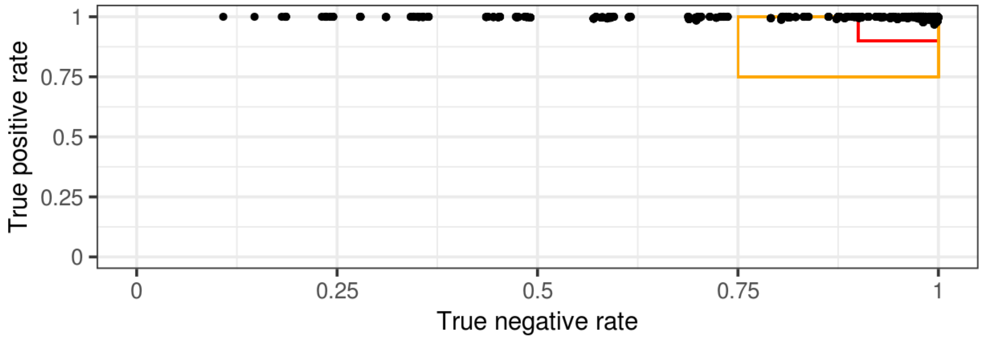


**Supplementary Figure S1.** Sensitivity (TP) vs specificity (TN) rates in all the runs performed with simulated data. The orange box in each facet represents “good” runs (TP and TN ≥ 0.75) while the red box represents “excellent” runs (TP and TN ≥ 0.9).

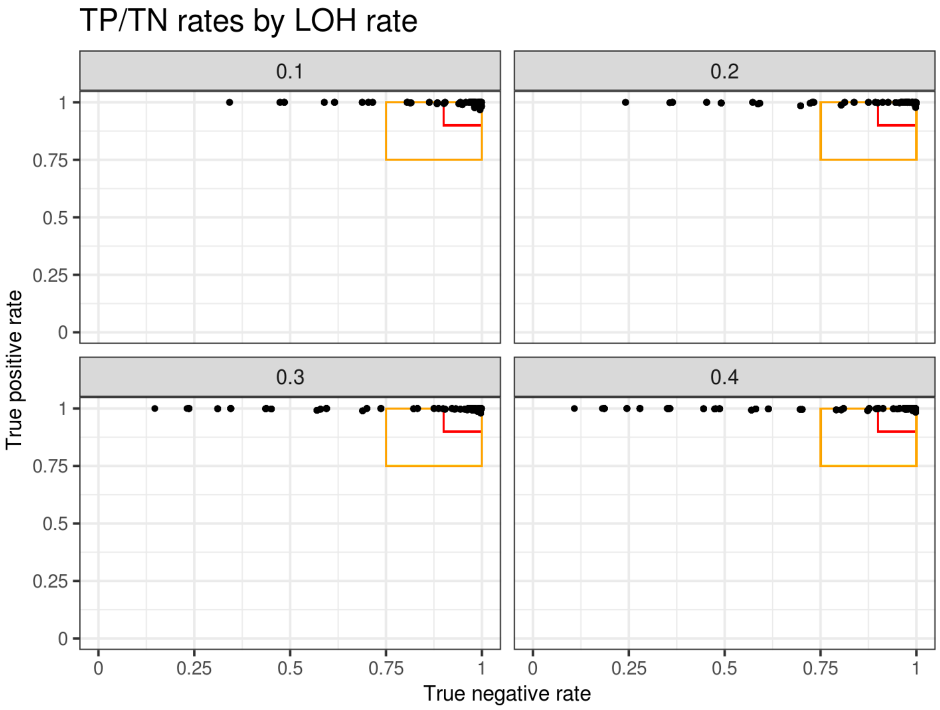


**Supplementary Figure S2.** Sensitivity (TP) vs specificity (TN) rates in all the runs performed with simulated data, subdivided in facets corresponding to the simulated rate of LOH (10%, 20%, 30%, 40%). The orange box in each facet represents “good” runs (TP and TN ≥ 0.75) while the red box represents “excellent” runs (TP and TN ≥ 0.9).

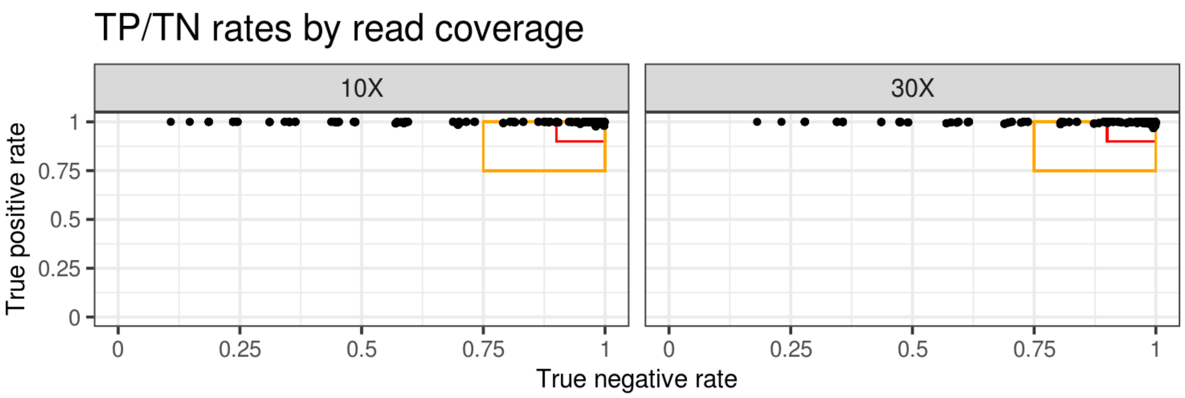


**Supplementary Figure S3.** Sensitivity (TP) vs specificity (TN) rates in all the runs performed with simulated data, subdivided in facets corresponding to the simulated sequencing coverage (10X and 30X). The orange box in each facet represents “good” runs (TP and TN ≥ 0.75) while the red box represents “excellent” runs (TP and TN ≥ 0.9).

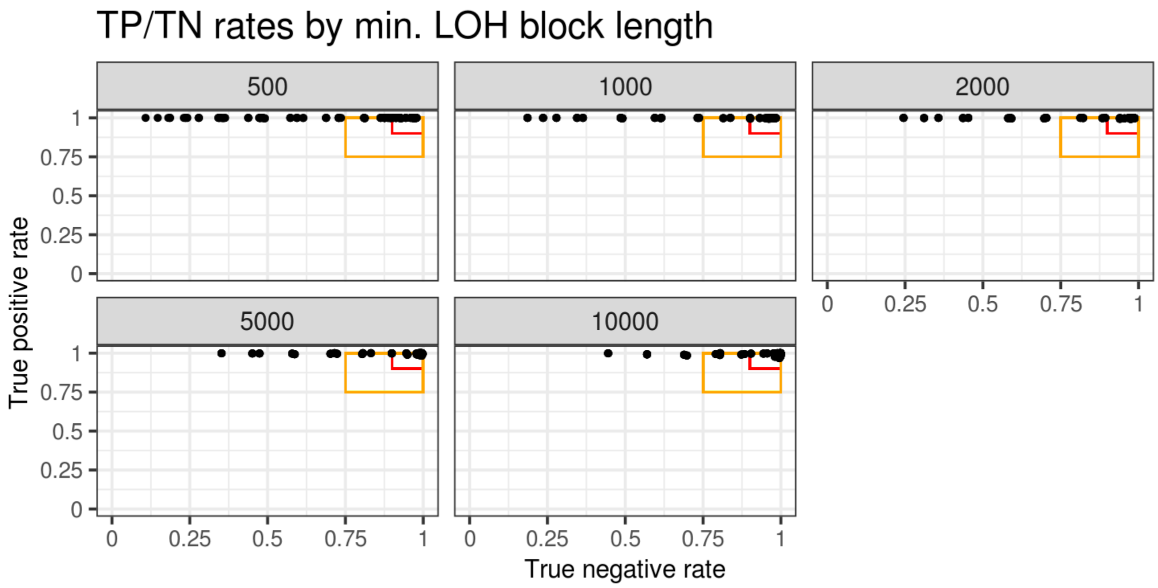


**Supplementary Figure S4.** Sensitivity (TP) vs specificity (TN) rates in all the runs performed with simulated data, subdivided in facets corresponding to the minimum tolerated length of the inferred LOH blocks (100 bp and 1000 bp). The orange box in each facet represents “good” runs (TP and TN ≥ 0.75) while the red box represents “excellent” runs (TP and TN ≥ 0.9).

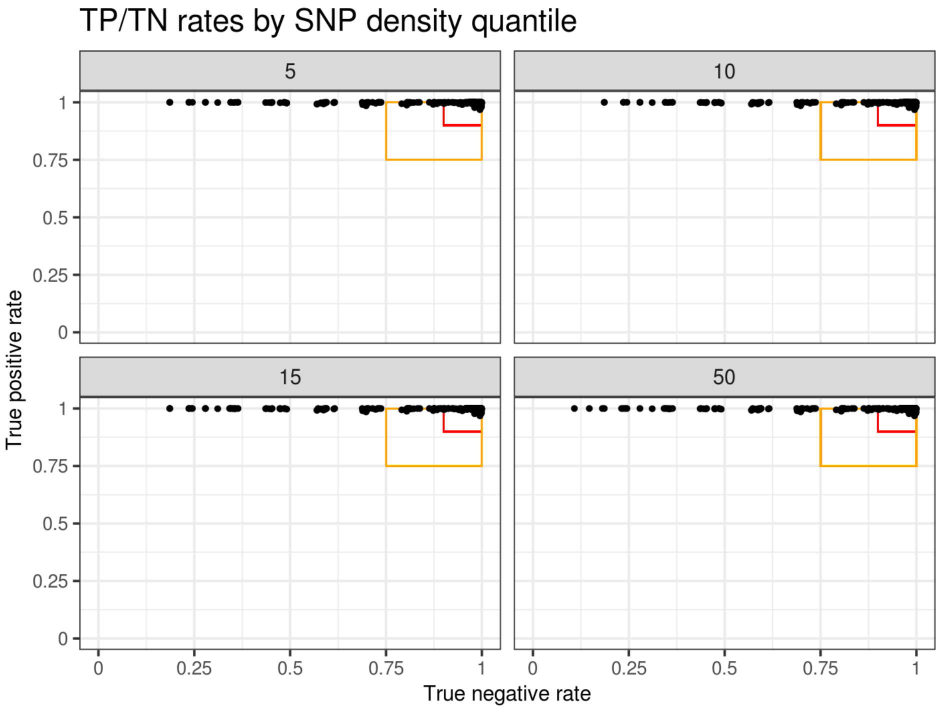


**Supplementary Figure S5.** Sensitivity (TP) vs specificity (TN) rates in all the runs performed with simulated data, subdivided in facets corresponding to the SNP density quantile selected (5, 10, 15, 50). The orange box in each facet represents “good” runs (TP and TN ≥ 0.75) while the red box represents “excellent” runs (TP and TN ≥ 0.9).

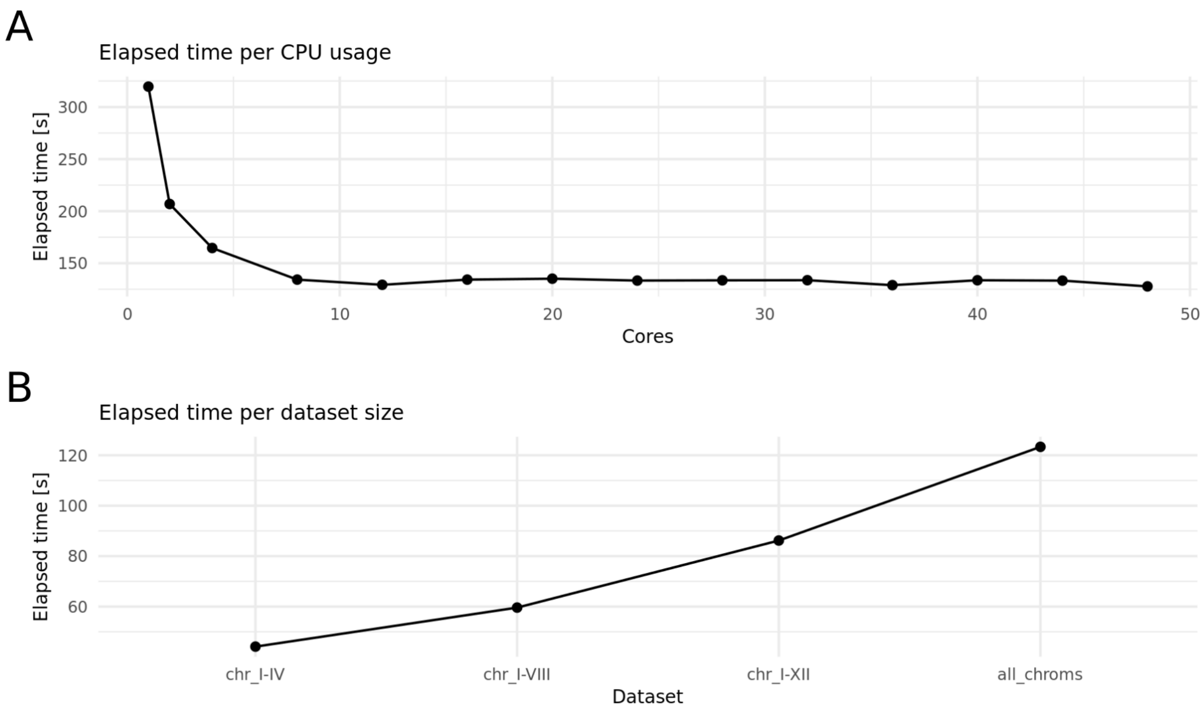


**Supplementary Figure S6**. Elapsed time measured in seconds (s) for each run performed with **A**) different number of CPUs (“Cores”), and **B**) different input data sizes (“Dataset”). Each black dot represents a different run.

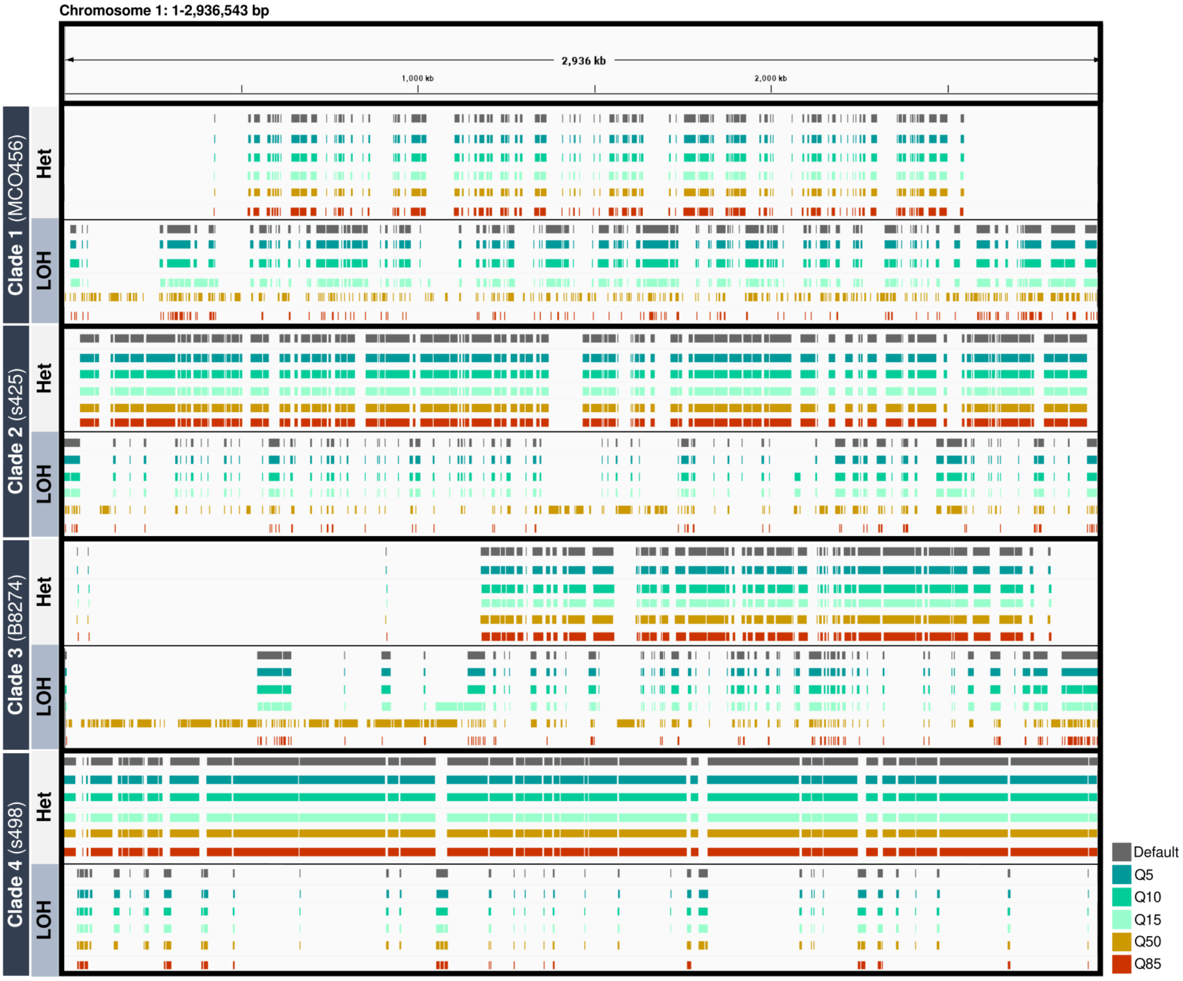


**Supplementary Figure S7.** Summary of LOH and heterozygous blocks along chromosome 1 of one representative *C. orthopsilosis* hybrid from each clade. Different coloured tracks represent different SNP density parameters used to infer LOH according to the results of the JLOH stats module.

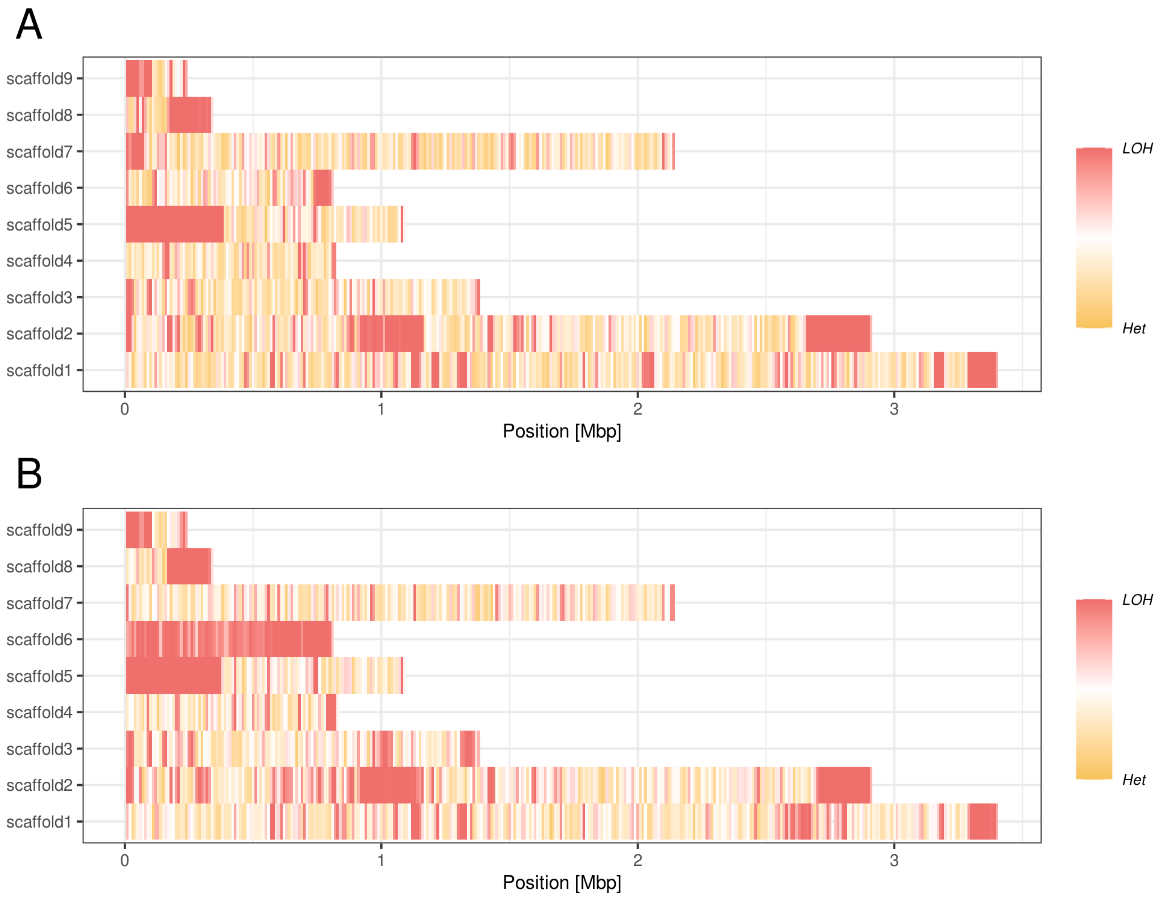


**Supplementary Figure S8.** LOH patterns displayed by the *C. metapsilosis* BP57 and PL448 strains. Color indicates LOH propensity from fully heterozygous (yellow) to fully homozygous (red). **A)** BP57 results; **B)** PL448 results.
