## Supplementary File description for "JLOH: Inferring Loss of Heterozygosity Blocks from Sequencing Data"

### Supplementary Files description

**Supplementary File S1.** Accession name and SRA code of the public hybrid yeast sequencing data.

**Supplementary File S2.** Summary of results obtained from JLOH stats and extract modules on *C. orthopsilosis* hybrid strains.

**Supplementary File S3.** True positives (TP), false positives (FP), true negatives (TN), false negatives (FN), precision, recall, specificity and sensitivity retrieved in each of the runs performed with simulated data. Run details are described in the first columns, in terms of SNPs/kbp, minimum length (bp), sequencing coverage (X), genome divergence (%), and LOH rate (%). The SNPs/kbp column indicates dash-separated SNP density thresholds for heterozygous and homozygous SNPs, e.g. “10-2”.

**Supplementary File S4.** Number of runs classified as basic (TP and TN ≥ 0.5), good (TP and TN ≥ 0.75), and excellent (TP and TN ≥ 0.9). Runs are stratified by parameter (SNPs/kbp, length, coverage) or by genomic property (divergence, LOH rate).

**Supplementary File S5.** Summary of shared LOH blocks between samples belonging to all clades and between samples belonging to the same clade obtained by running JLOH extract using the different --snp-per-kbp parameters obtained in the JLOH stats module.
